# HPAI H5N1 risk in Australia: a model for the prediction of poultry outbreaks

**DOI:** 10.64898/2026.07.27.740638

**Authors:** Pan Zhang, Samsung Lim, Ashley Quigley, C Raina MacIntyre

## Abstract

The panzootic highly pathogenic avian influenza (HPAI) H5N1 virus has now been detected on the Australian mainland, with incursions from the sub-Antarctic region posing an increasing threat to domestic wildlife and poultry populations. Our study aimed to predict the risk of HPAI H5N1 poultry outbreaks across Australia at the local government area (LGA) level using a range of influential risk factors. We first used a Maximum Entropy (MaxEnt) model to estimate the environmental suitability for HPAI H5N1 occurrence across Australia. The resulting suitability layer was then integrated with five additional predictor layers, including abundance data for two Southern Ocean wild birds, one of which has introduced HPAI H5N1 into Australia; abundance data for 28 native Australian wild birds; native bird flyways across Australia; Australian chicken density; and poultry farm density. The six layers were aggregated and averaged to generate an HPAI H5N1 risk map for poultry outbreaks across Australian LGAs. Although most incursions have occurred in Western Australia (WA) and South Australia (SA), we identified New South Wales (NSW) and Victoria (VIC) as having the highest predicted risk of HPAI H5N1 poultry outbreaks. Additional high-risk areas were identified in WA, SA, and Tasmania (TAS). In contrast, the Northern Territory (NT) and large parts of Queensland (QLD), WA, and SA were predicted to be at low risk. These findings provide a spatially explicit framework to support targeted surveillance, preparedness, and biosecurity measures aimed at mitigating the impact of future HPAI H5N1 outbreaks in Australian poultry.

## Introduction

### Current situation in Australia

The panzootic highly pathogenic avian influenza (HPAI) H5N1 virus of clade 2.3.4.4b has now been discovered in mainland Australia [1]. Since its emergence in late 2020 [2], Australia has remained free of incursions of this lineage [3]. However, this status changed after the virus was detected on Australia’s sub-Antarctic territory, Heard Island, in November 2025 [4], and subsequently on mainland Australia in June 2026 [1].

The first mainland case was confirmed on 20 June 2026 in a brown skua (*Stercorarius antarcticus*), a migratory seabird from the sub-Antarctic region, at Cape Le Grand National Park near Esperance, Western Australia (WA) [5]. Since this initial detection, reports of HPAI H5N1 cases have persisted across various states, including South Australia (SA) [6] and New South Wales (NSW) [7,8]. As of 20 July 2026, there have been a total of 17 confirmed cases of HPAI H5N1 in wild avian populations throughout Australia [9]. Of these, most cases involved giant petrels (*Macronectes*), which are large migratory seabirds of the Southern Ocean. One instance was reported in a Greater Crested Tern (*Thalasseus bergii*), representing the first identification of HPAI H5N1 in a native Australian wild bird species [8]. Currently, cases of HPAI H5N1 clade 2.3.4.4b have not been reported among other populations, including other native wild birds, marine mammals, domestic mammals, poultry, or humans within Australia.

### Viral evolution and animal health risks associated with HPAI H5N1 introduction

The introduction of the panzootic clade 2.3.4.4b HPAI H5N1virus into new continents has been accompanied by extensive genomic reassortment with locally circulating low-pathogenic avian influenza (LPAI) viruses [10]. These reassortment events have generated numerous novel genotypes, facilitating adaptation to a broader range of hosts and contributing to widespread outbreaks in both avian and mammalian populations [10,11,12]. In North America, the introduction of Eurasian clade 2.3.4.4b HPAI H5N1 via migratory birds in late 2021 was followed by reassortment with endemic LPAI viruses, resulting in the emergence of B and D genotype groups [13,14]. The B3.13 genotype was associated with the unprecedented outbreak in U.S. dairy cattle since March 2024, whereas the D1.1 genotype later became prevalent in wild birds and was responsible for multiple poultry outbreaks before being detected in cattle [14].

A similar pattern was observed in South America following the introduction of clade 2.3.4.4b HPAI H5N1 in late 2022 [15]. Believed to have originated from the North American B3.2 genotype, the virus caused extensive outbreaks in poultry and colonial seabirds. Subsequently, a divergent viral lineage emerged in marine mammals, resulting in mass mortality events among pinniped species, including southern elephant seals (*Mirounga leonina*), South American sea lions (*Otaria flavescens*), and South American fur seals (*Arctocephalus australis*) [15].

The detection of HPAI H5N1 clade 2.3.4.4b in Australia therefore raises concerns regarding its potential impact on native animal health [16]. In addition to the direct threat posed to poultry populations, the virus may affect a wide range of native wild bird and mammalian species. Australia’s history of hosting diverse LPAI viruses [17] may enhance the likelihood of reassortment with the introduced HPAI H5N1 virus, which could potentially modify the AIV community in Australia and generate novel genotypes with expanded host adaptation [16].

### Study aim

Under this threat, the immediate impact of HPAI H5N1 is likely to be greatest in poultry populations. In the U.S., the virus caused losses of approximately 108 million birds from commercial and backyard poultry flocks between early 2022 and late 2024, with outbreaks reported across nearly all states [18]. In Europe, HPAI H5N1 resulted in the culling of, or infection among, an estimated more than 120 million domestic poultry birds between 2020 and 2025 [19]. Peak activity occurred during 2021-2022, when over 2,500 poultry outbreaks were reported across more than 30 European countries [19]. The threat of HPAI H5N1 outbreaks in Australian poultry is expected to escalate as the virus spreads among wild birds, widening the range of flyways for transmission to poultry. Therefore, this study aimed to predict the risk of HPAI H5N1 poultry outbreaks across Australia at the local government area (LGA) level using related predictors. By identifying areas at elevated risk, our model can support targeted surveillance, prevention, and biosecurity measures to reduce the potential impact of future outbreaks in Australian poultry.

## Materials and methods

Our study incorporated a range of variables to predict the risk of HPAI H5N1 poultry outbreaks in Australia. The following sections provide details on the data sources, modelling methodologies, and data processing and analytical procedures used in this study.

### Data collection and preprocessing procedures

#### Study area

The study area encompassed Australia, including mainland Australia and Tasmania, with a spatial extent of 112.89-159.14°E and 9.12-43.79°S.

#### HPAI H5N1 cases

We used 17 H5N1 cases reported in Australia as of July 20, 2026. Case data were retrieved from publicly available sources, including news articles and government reports [5,6,7,8,9]. The geographic coordinates (latitude and longitude) of the reported locations were estimated using Google Maps.

#### Bioclimatic variables

WorldClim version 2.1 [20,21] is a broadly used platform that provides high-resolution (approximately 1 km^2^) global climate data based on historical observations collected between 1970 and 2000. The dataset had been extensively applied in ecological modelling studies, including investigations of species distribution and disease risk [22]. We downloaded historical monthly climate data for the period 2020-2024 at a spatial resolution of 2.5 arcminutes (approximately 21km^2^ at the equator) [20,21]. To assess the immediate risk of HPAI H5N1 poultry outbreaks in Australia as of July 2026, we selected precipitation and minimum temperature data for July 2024 for inclusion in the prediction model. WorldClim distributes these climate data in raster (tiff) format with global coverage.

#### Wetlands

Our study incorporated Ramsar wetlands into the prediction model because they represent wetlands of international importance and provide critical habitat for waterbirds [23], which may contribute to HPAI H5N1 transmission dynamics. Although Australia contains numerous nationally important wetlands, Ramsar wetlands offer a standardized and internationally recognized framework for identifying ecologically significant sites. These wetlands encompass a wide range of natural or human-made habitats, including rivers, lakes, swamps, coral reefs, and other aquatic ecosystems [24]. Australia currently hosts 67 Ramsar wetlands, covering over 8.3 million hectares [25].

One of the key criteria for Ramsar designation is the regular support of at least 20,000 waterbirds, highlighting the importance of these sites as major waterbird habitats [24]. Ramsar Wetlands serve as important habitats for waterfowl, shorebirds, and migratory species, which can carry avian influenza viruses (AIVs) and move between regions [26]. By incorporating Ramsar wetlands into the model, we aimed to identify areas with a high likelihood of HPAI H5N1 introduction and circulation within wild avian populations. The spatial shapefile of Australian Ramsar Wetlands was obtained from the ArcGIS Pro 3.5 online portal [27] maintained by the Australian Government Department of Climate Change, Energy, the Environment and Water (DCCEEW). To quantify spatial proximity to Ramsar Wetlands, we generated a Euclidean distance raster using ArcGIS Pro 3.5 [28].

#### Wild bird abundance

In this study, the HPAI H5N1 risk of outbreaks in Australian poultry was assessed through a transmission pathway involving migratory seabirds from the Southern Ocean, native Australian wild birds, and domestic poultry. Specifically, we hypothesized that HPAI H5N1 viruses introduced by migratory seabirds may be subsequently transmitted to native wild birds and ultimately spread to poultry populations.

To represent these transmission pathways, two sets of wild bird abundance data were obtained from the eBird Status and Trends database [29]. The first dataset focused on two representative Southern Ocean migratory species: the Northern Giant Petrel (*Macronectes halli*) and the Fairy Prion (*Pachyptila turtur*). The Northern Giant Petrel was included because it accounted for the majority of the 17 HPAI H5N1 detections reported in Australia as of 20 July 2026. Both species are closely associated with southern Australian waters and offshore islands and were selected as potential carriers of the virus into Australia.

The second dataset comprises 28 native Australian wild bird species representing 11 taxonomic orders (**Supplementary Table S1**). Among these, species within the orders *Anseriformes* and *Charadriiformes* are recognized as important reservoirs and vectors of HPAI H5N1 viruses [30] and may contribute to virus transmission to domestic poultry. In addition, species belonging to the order *Accipitriformes* were considered important components of the transmission pathway because, as predatory raptors and scavengers, they are likely to become infected through contact with or consumption of infected bird carcasses [31].

The eBird Status and Trends database [29] provides model-based estimates of bird abundance at an approximate spatial resolution of 14km x 14km in raster format (tiff) rather than raw observation counts [32]. The mean abundance raster layer for each bird dataset (i.e., the two Southern Ocean migratory seabird species and the 28 native wild bird species) was calculated and created separately in ArcGIS Pro 3.5 for subsequent modelling.

#### Flyways across Australia

Using the mean abundance raster derived from the 28 native Australian wild bird species as described above, we created a flyway raster layer to represent potential movement corridors across Australia. Specifically, bird abundance was expanded using a 100-km radius neighbourhood, and a smoothed raster surface was generated in ArcGIS Pro 3.5. This preprocessing step was undertaken to account for the mobility of wild bird populations and to capture potential dispersal pathways beyond areas of observed abundance. By incorporating these potential movement corridors, the flyway layer was designed to maximize the spatial extent of areas with a likelihood of contact between native Australian wild birds and poultry populations. This approach aimed to capture potential HPAI H5N1 transmission pathways.

#### Chicken density

Chicken density data for the reference year 2020 were obtained from Version 4 of the Gridded Livestock of the World (GLW-4) dataset [33] maintained by the Food and Agriculture Organization (FAO). The dataset provides global livestock density estimates at a spatial resolution of 5 arcminutes (approximately 10km at the equator).

#### Poultry farms

Locations of poultry farms in Australia were obtained from the Farm Transparency Project database [34]. The farm location coordinates were imported into ArcGIS Pro 3.5, where a poultry farm density layer was generated to represent the spatial distribution and concentration of poultry production facilities across the study area.

Before modelling, we cropped and masked all raster layers in ArcGIS Pro 3.5 to the spatial extent of the study area.

### Modelling methods

In this study, prediction modelling was conducted using Maximum Entropy (MaxEnt) and Geoinformation System (GIS)-based risk integration. We first used the MaxEnt model to estimate the probability of environmental suitability for HPAI H5N1 using 17 detections and three environmental predictors. Subsequently, the MaxEnt-derived HPAI H5N1 environmental suitability layer was integrated in ArcGIS Pro 3.5 with five additional raster layers: (1) the mean abundance raster of two Southern Ocean migratory seabirds, (2) the mean abundance raster of 28 native Australian wild bird species, (3) the flyway layer, (4) the Australian chicken density layer, and (5) the Australian poultry farm density layer. We summed the six layers and divided the result by six to generate a composite mean raster representing the risk of HPAI H5N1 poultry outbreaks.

This predictive modelling approach took into consideration the risk of HPAI H5N1 introduction by Southern Ocean migratory seabirds, the environmental suitability of the virus across Australia, the potential for HPAI H5N1 transmission from Southern Ocean Migratory seabirds to native Australian wild bird species, the subsequent dispersal of the virus through local wild bird movement, and the risk of transmission to domestic poultry populations.

The MaxEnt model, widely used in ecological studies, identifies and predicts the probability of species occurrence under a given set of environmental conditions [35,36,37]. Building on this concept, researchers have increasingly applied the MaxEnt model to disease risk assessment, particularly for AIVs [32,38]. Using presence-only data, the model predicts areas of environmental suitability and potential disease occurrence under specific environmental conditions.

As only 17 HPAI H5N1 cases had been reported in Australia as of 20 July 2026 [9], we incorporated a limited set of three environmental variables into MaxEnt version 3.4.3 [39]: Euclidean distance to Ramsar Wetlands, July 2024 precipitation, and July 2024 minimum temperature. Because the reported HPAI H5N1 cases were limited and only concentrated along the southern coastline of Australia, and large areas of inland Australia are characterized by arid and semi-arid conditions that are unlikely to support HPAI H5N1 persistence, we reduced the number of background points from the default value of 10,000 to 5,000. This adjustment aimed to improve the model’s ability to identify environmental patterns associated with limited occurrence records. We subsequently used the calibrated MaxEnt model to estimate the environmental suitability of HPAI H5N1 across mainland Australia. The resulting MaxEnt suitability values range from 0 to 1, with higher values indicating greater environmental suitability for HPAI H5N1 occurrence. Five replicate MaxEnt runs were performed, and the mean prediction across all five replicates was selected for subsequent analyses.

All other raster layers were normalized in ArcGIS Pro 3.5 to a common scale ranging from 0 to 1. Finally, the resulting risk raster was integrated with Australian Local Government Area (LGA) boundaries in ArcGIS Pro 3.5 to visualise the spatial distribution of outbreak risk at the local government level. The risk values were classified into five categories and labelled as Very Low, Low, Moderate, High, and Very High to facilitate the interpretation.

We also developed an interactive map that displays all predictor-variable layers and the predicted HPAI H5N1 poultry outbreak risk raster at the following URL: https://ui.epiwatch.app/avian_map/

## Results

### HPAI H5N1 environmental suitability results based on MaxEnt modelling

The MaxEnt model achieved a mean AUC of 0.965 across five replicate runs, indicating excellent performance. **Figure 1** exhibits the average MaxEnt-modelled prediction for HPAI H5N1 environmental suitability based on the 17 reported occurrence records and three environmental variables. The model identified the south-west corner of WA as the region with the highest environmental suitability for HPAI H5N1 occurrence. Other areas with high suitability included western and northern Tasmania (TAS), eastern Queensland (QLD), and the southern coastline of Victoria (VIC), with predicted values ranging from 0.85 to 1. Areas of moderate suitability (0.46-0.77) occurred across parts of the Northern Territory (NT), northern and southern WA, eastern coastal regions of Australia, the southern coastal zones of SA and VIC, as well as the coastal regions of TAS. In contrast, most inland regions of Australia exhibited very low environmental suitability for HPAI H5N1 occurrence according to the MaxEnt model. Additionally, portions of central NSW and VIC displayed suitability values ranging from 0.31 to 0.38.

**Figure 1.**
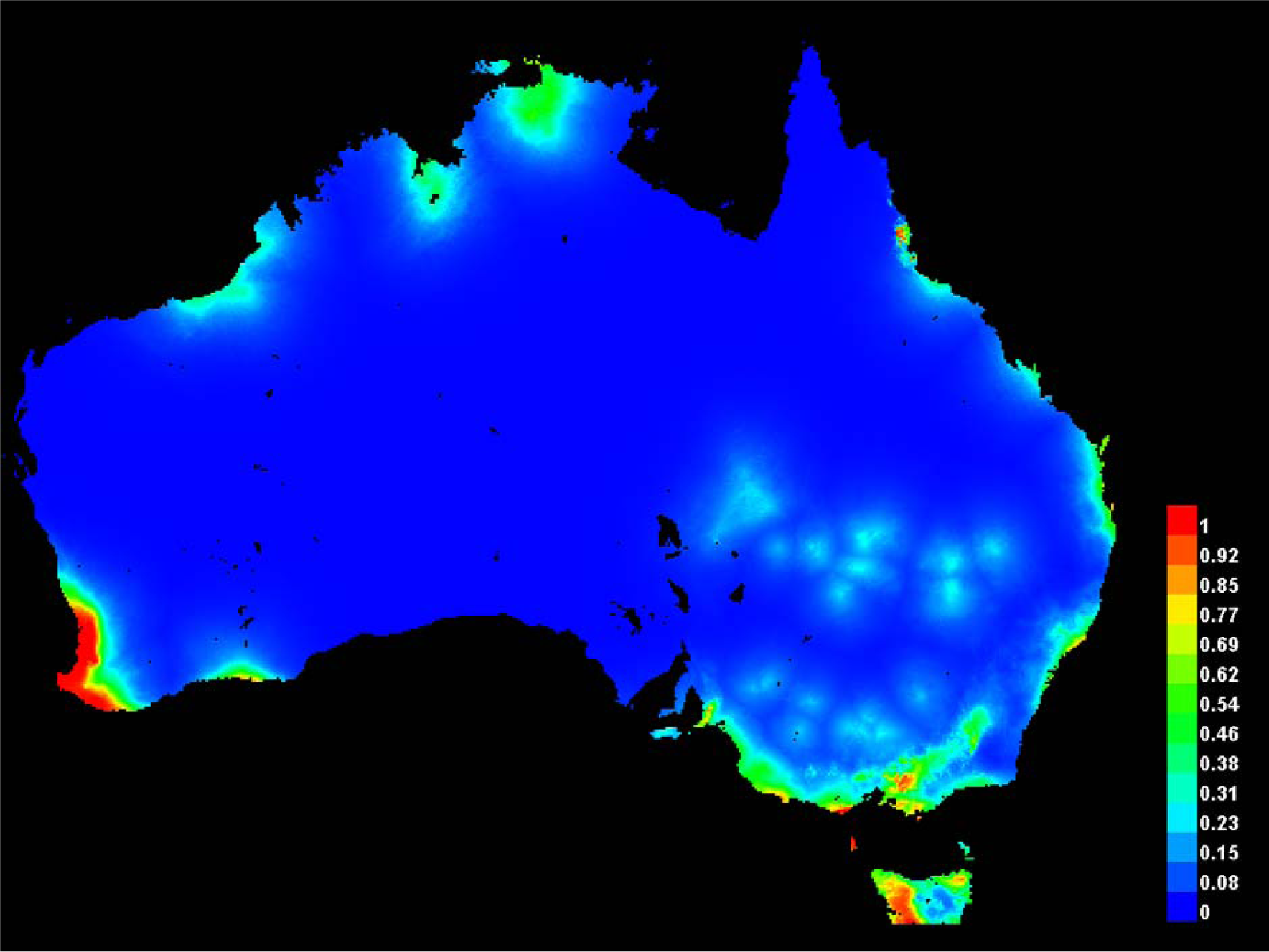
MaxEnt-predicted HPAI H5N1 environmental suitability across Australia. The model was fitted using 17 presence records reported as of 20 July 2026 and included three environmental predictors: Euclidean distance to Ramsar Wetlands, July 2024 precipitation, and July 2024 minimum temperature. Suitability values range from 0 to 1, where values closer to 1 indicate a higher probability of environmental suitability for HPAI H5N1 occurrence.

Figure 2. presents the response curves for the three environmental variables, and **Figure 3** presents the jackknife evaluation of variable importance. Among the environmental variables included in the model, Jackknife analysis (**Figure 3**) revealed that July 2024 precipitation contributed the most to the model predictions. Models trained using July 2024 precipitation alone achieved the highest gain, while omitting this variable resulted in a marked reduction in model performance, suggesting that precipitation contained substantial unique information relevant to HPAI H5N1 environmental suitability.

**Figure 2.**
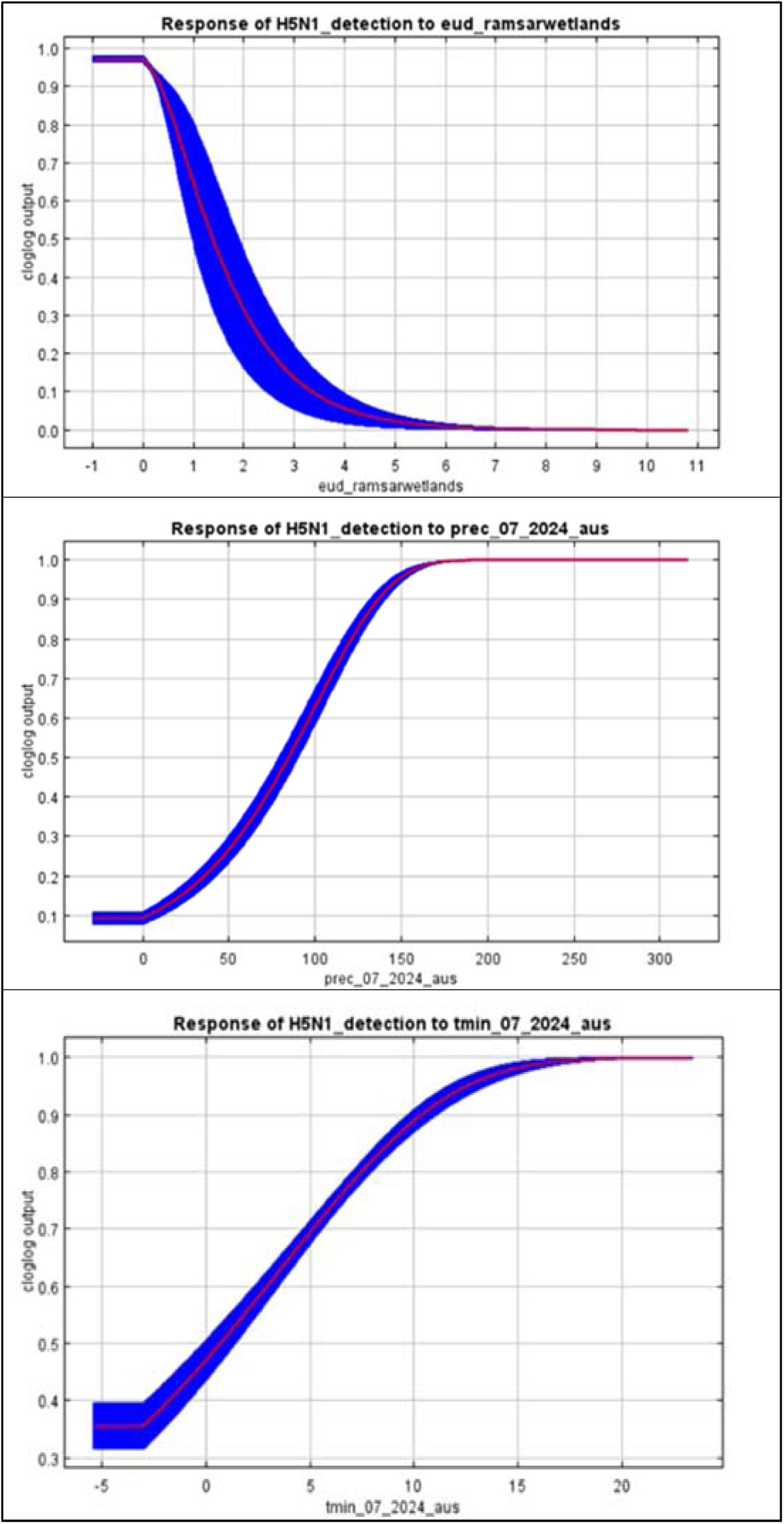
Response curves of three environmental variables.

**Figure 3.**
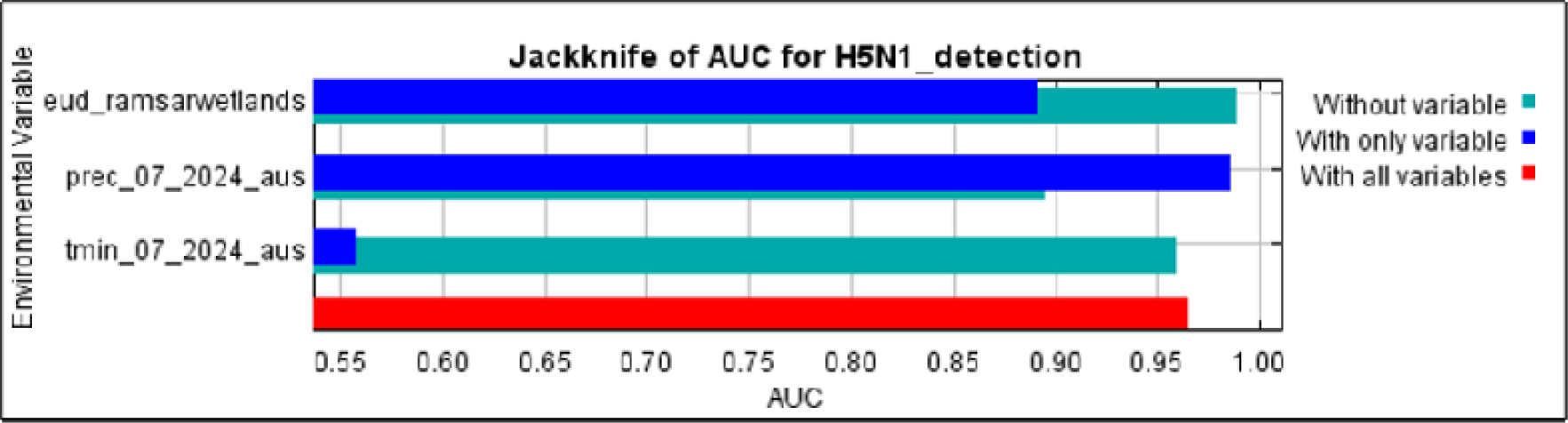
Jackknife evaluation of the relative importance of three environmental variables.

### HPAI H5N1 risk prediction of poultry outbreaks

Our modelling prediction, derived from averaging the six risk-factor layers, identified potential risk areas for HPAI H5N1 poultry outbreaks across southern Australia and along the east coast (**Figure 4**). The highest predicted risk areas were concentrated in NSW and VIC.

**Figure 4.**
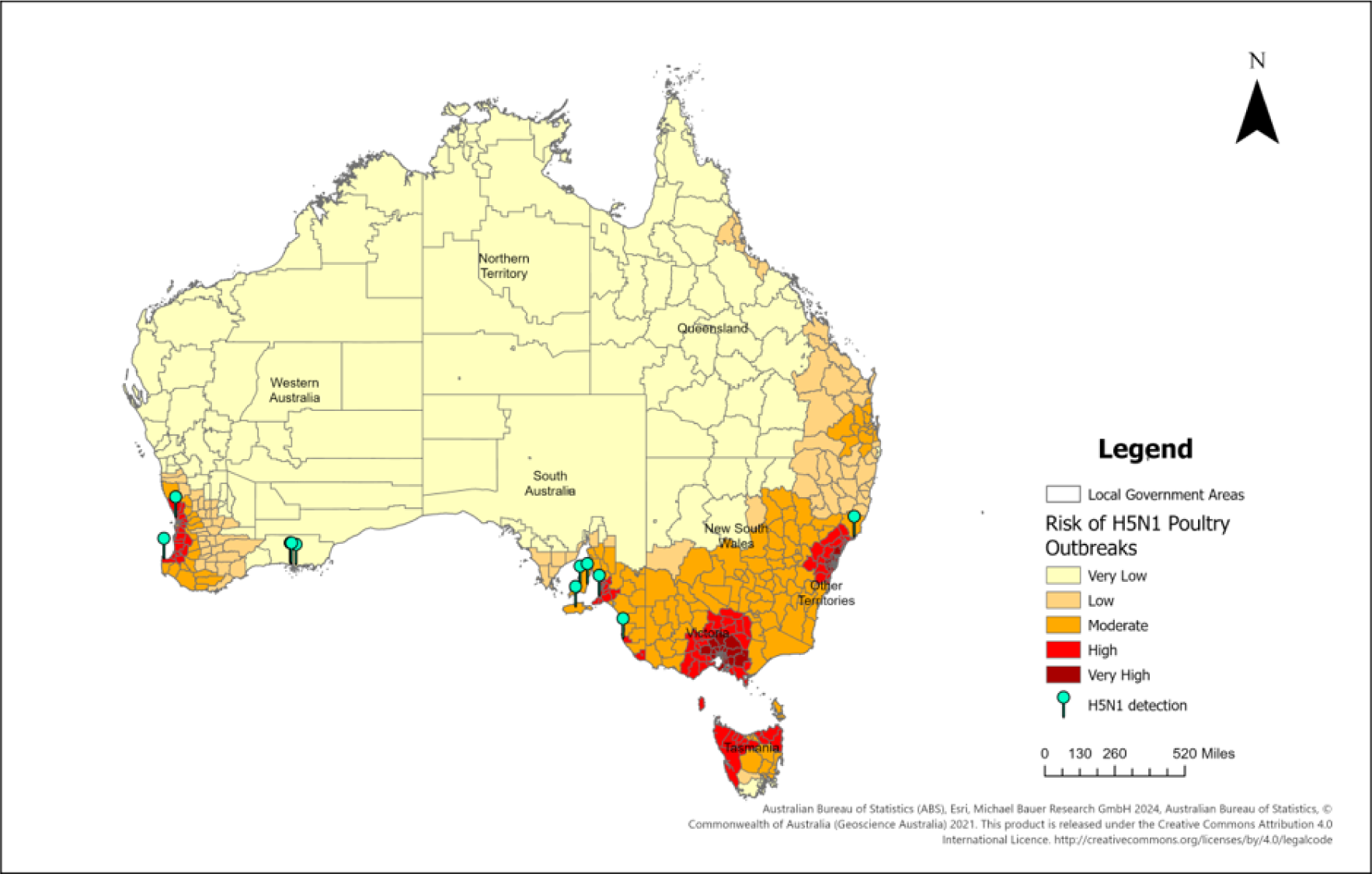
Predicted spatial risk of HPAI H5N1 poultry outbreaks across Australian local government areas.

In NSW (**Figure 5**), the LGAs classified as very high risk included Central Coast, Hornsby, Northern Beaches, The Hills, Ku-ring-Gai, Parramatta, Penrith, Cumberland, Fairfield, Liverpool, Canterbury-Bankstown, Camden, Campbelltown, and Sutherland. Notably, NSW had already reported a case of HPAI H5N1 in a giant petrel at Hawks Nest (Central Coast-Port Stephens corridor) in early July 2026 [7,8]. Although this case occurred within an LGA classified as having moderate risk, it was located adjacent to several LGAs identified as high risk in our modelling prediction, including Port Stephens, Maitland, Newcastle, Cessnock, and Hawkesbury. These findings suggest an elevated risk of H5N1 poultry outbreaks in NSW associated with the already reported occurrence in migratory bird species and highlight the need for enhanced surveillance and biosecurity measures in neighbouring regions.

**Figure 5.**
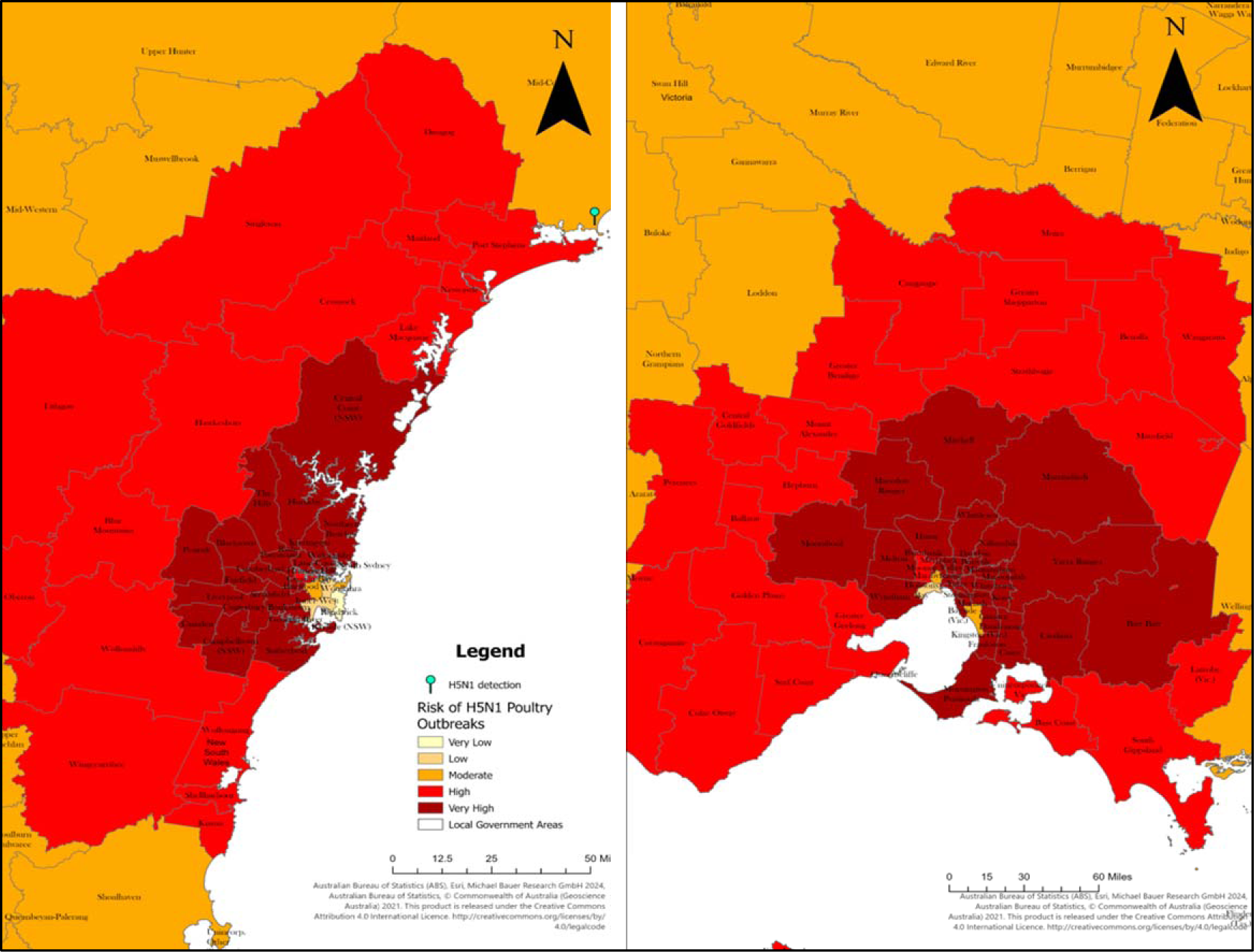
Enlarged view of predicted very high-risk areas for HPAI H5N1 poultry outbreaks in New South Wales (NSW) (left) and Victoria (VIC) (right)

Our modelling also identified several very high-risk LGAs within VIC (**Figure 5**), including Yarra Ranges, Knox, Wyndham, Nillumbik, Manningham, Darebin, Baw Baw, Hume, Whittlesea, Melton, Mitchell, Moorabool, Murrindindi, Cardinia, and Casey. Surrounding these areas, many LGAs were classified as high risk, including Latrobe, Surf Coast, Golden Plains, and Greater Bendigo.

In addition, southern SA, the south-west corner of WA, and northern TAS also contained areas predicted to be at high risk of H5N1 poultry outbreaks. In contrast, the NT exhibited predominantly very low risk for poultry outbreaks. Similarly, large parts of WA, SA, and QLD were classified as having low to very low risk.

## Discussion

The detection of HPAI H5N1 in a Greater Crested Tern (*Thalasseus bergii*), a native Australian wild bird species, in South Australia on 10 July 2025 represents an important epidemiological development [6]. While most detections to date have been reported in migratory seabirds originating outside the mainland from the Southern Ocean, the occurrence of the infection in a native Australian wild bird species suggests that the virus may be expanding beyond its initial introduction pathway. This raises concerns that HPAI H5N1 could become more widely distributed through movements and interactions among local wild bird populations, thereby increasing opportunities for transmission to susceptible poultry populations. Consequently, the risk of HPAI H5N1 poultry outbreaks may escalate rapidly, particularly in regions where environmental conditions, wild bird activity, and poultry production systems favour viral persistence and spread.

Although as of 20 July 2026, no other states have reported HPAI H5N1 detections in local wild bird species, the implications extend beyond South Australia, where the detection in a Greater Crested Tern (*Thalasseus bergii*) was reported. NSW and VIC warrant particular attention, as both states host extensive poultry production systems [34] and have experienced a disproportionate burden of avian influenza outbreaks in poultry in the past [40]. Notably, during 2024 and 2025, HPAI H7 viruses caused substantial increases in poultry outbreaks across both states [41]. The identification of Hawkesbury, NSW, and Golden Plains, VIC, as high-risk areas within our model is consistent with historical patterns, suggesting that these regions may continue to foster the emergence of avian influenza, such as HPAI H5N1, in poultry populations. While VIC has not reported HPAI H5N1 detections to date, and SA remains the only state with a confirmed detection in a native wild bird, the concentration of predicted very high-risk areas for poultry outbreaks in NSW and VIC underscores the necessity for heightened preparedness. The stringent protection of poultry from exposure to native wild bird species in these two states constitutes a vital biosecurity measure.

Compared with NSW and VIC, WA, SA, and TAS generally exhibit lower densities of poultry farms [34]. Nonetheless, the high-risk areas identified within these states by our model may be attributable to an increased probability of HPAI H5N1 virus introduction via migratory seabirds associated with the Southern Ocean. Numerous migratory seabird species frequent southern Australian waters [42], indicating multiple potential entry points for viral incursion along the southern coastline, including regions such as WA, SA, VIC, and TAS. Such introductions could heighten the likelihood of spillover into endemic wild bird populations, consequently amplifying the risk of subsequent transmission to domestic poultry.

Furthermore, the MaxEnt model predicted relatively high environmental suitability for the establishment of HPAI H5N1 in several regions within these states following viral introduction, which aligns with the spatial distribution of areas identified as high-risk for poultry outbreaks. Therefore, enhanced surveillance of HPAI H5N1 in external migratory seabirds and the implementation of strengthened biosecurity measures to prevent viral spillover into local wild bird species, particularly in WA, SA, VIC, and TAS, may be essential for mitigating the rapid emergence of poultry outbreaks.

The comparatively low predicted risk of poultry outbreaks across much of northern Australia, including northern WA, the NT, and parts of QLD, should be interpreted in the context of both the model assumptions and regional poultry production patterns. Specifically, the current modelling framework primarily considered the introduction of HPAI H5N1 via migratory seabirds from the Southern Ocean and its subsequent spread from the southern coast of Australia. As a result, it did not explicitly account for potential viral incursions through northern migratory pathways, such as the East Asian-Australasian Flyway (EAAF), despite the possibility of such introductions [43]. This modelling design may have resulted in an underestimation of HPAI H5N1 introduction risk in northern regions. However, the overall risk of HPAI H5N1 incursion into northern Australia is considered low, likely due to the biogeographic barrier imposed by Wallace’s Line [44]. Although the MaxEnt model identifies some areas in northern Australia as environmentally suitable for HPAI H5N1, the relatively low density of poultry farms and chickens across these regions may reduce their opportunities for viral exposure, resulting in a lower risk of poultry outbreaks. Nevertheless, as additional HPAI H5N1 detections occur and the distribution of the virus changes, the risk profile may shift, highlighting the need for ongoing surveillance and regular updates of risk assessments.

This study has several limitations. First, the limited number of HPAI H5N1 occurrence records constrained the range of environmental variables that could be incorporated into the MaxEnt model. This may have reduced its ability to accurately characterise true disease distribution patterns under different environmental conditions. The model calculated only precipitation, minimum temperature, and the distance to Ramsar wetlands as predictors of HPAI H5N1 occurrence. Although these variables are likely to influence viral survival and persistence, their specific associations with HPAI H5N1 were not quantitatively evaluated before inclusion in the model. Furthermore, all raster layers were masked to a spatial resolution of approximately 300m. This resolution was selected to ensure accurate calculations and consistency across all predictor raster layers. However, overlaying the final poultry outbreak risk raster with LGA boundaries may have resulted in missing values by default for a small number of LGAs, particularly those with very small geographic extents. Nevertheless, the extent of this issue was minimal. Despite these limitations, the high-risk areas identified by our model are generally consistent with the patterns of historical poultry outbreaks in Australia, supporting the robustness of the modelling framework and the validity of the findings. This provides data to inform and prioritise biosecurity measures for Australia.

## Ethics

The study used open-source data without identifying information. As such, ethics approval was not required.

## APPENDIX

**Supplementary Table S1.** List of 28 native Australian wild bird species whose abundance raster layers were included in the prediction model.

| ORDER | SPECIES |
| --- | --- |
| **ANSERIFORMES** | 1.Australian Shelduck<br>2.Australasian Shoveler<br>3.Pink-Eared Duck<br>4.Musk Duck<br>5.Pacific Black Duck<br>6.Black Swan<br>7.Magpie Goose |
| **CHARADRIIFORMES** | 1.Australian Fairy Tern<br>2.Bar-Tailed Godwit<br>3.Red-Necked Stint<br>4.Far Eastern Curlew<br>5.Silver Gull<br>6.Masked Lapwing<br>7.Greater Crested Tern |
| **ACCIPITRIFORMES** | 1.Wedge-Tailed Eagle<br>2.Swamp Harrier<br>3.Whistling Kite |
| **CICONIIFORMES** | 1.Straw-Necked Ibis<br>2.White-Faced Heron<br>3.Australian White Ibis |
| **PASSERIFORMES** | 1.Australian Magpie<br>2.Australian Raven |
| **PSITTACIFORMES** | 1.Australian King-Parrot |
| **GRUIFORMES** | 1.Brolga |
| **GALLIFORMES** | 1.Australian Brush-Turkey |
| **COLUMBIFORMES** | 1.Crested Pigeon |
| **PROCELLARIIFORMES** | 1.Short-tailed Shearwater |
| **PSITTACIFORMES** | 1.Rainbow Lorikeet |

